# Profile analysis in listeners with sensorineural hearing loss

**DOI:** 10.1101/2023.11.30.569471

**Authors:** Daniel R. Guest, David A. Cameron, Douglas M. Schwarz, U-Cheng Leong, Laurel H. Carney

## Abstract

Many sounds contain spectral modulations at multiple scales, but much is still unknown about how such spectral features are represented in the auditory system. One behavioral task that provides insight into this question is profile analysis. In a typical profile-analysis task, listeners are asked to discriminate between a complex tone with equal-amplitude components and a complex tone with a single incremented component. Because listeners can perform profile analysis even when the overall sound level of the stimuli is randomized from interval to interval, this task is thought to be a useful index of relative processing of spectral shape, rather than just sensitivity to absolute level changes. Here, we measured profile analysis across the frequency range in a group of listeners that varied widely in their hearing status. We then modeled the resulting behavioral data by decoding responses to the stimuli from computational models of the auditory nerve and inferior colliculus. We found that both hearing loss at the target frequency and increases in the target frequency were associated with poorer profile-analysis thresholds, and that these results could both be explained as the result of corresponding changes in sensitivity of temporal modulation-sensitive cells at the level of the inferior colliculus. These results suggest that key features of profile-analysis may reflect the limits of central neural tuning to temporal modulations.

## I. INTRODUCTION

Many ecologically relevant audio signals contain rich and dynamic spectral modulations at multiple scales. For example, speech contains fine spectral details created by the periodicity of voicing, spectral peaks created by the resonances of the vocal tract (i.e., formants), and idiosyncratic patterns of spectral peaks and troughs imparted by the shape of the head and torso (i.e., head-related transfer functions). These spectral patterns are a key means by which information is transmitted in speech communication, but there are still major gaps in our understanding of how such patterns are coded by the auditory system, especially in challenging acoustic contexts, such as noisy and unpredictable backgrounds, or in impaired auditory systems. Additionally, it remains to be seen how these spectral patterns are coded in relative terms, rather than absolute terms, such that coding is robust to unpredictable changes in overall sound level (e.g., randomly varying levels in both the real world and in psychoacoustic experiments).

One relevant line of inquiry is profile analysis, which tests the ability of listeners to discriminate changes in the amplitude of a single component relative to that of its neighboring components in a complex tone, or of sets of components relative to others (Spiegel et al., 1981; Green, 1983; Green et al., 1983; Green and Mason, 1985; Bernstein and Green, 1987, 1988; Richards et al., 1988; Kidd et al., 1991; Gockel and Colonius, 1997; Gockel, 1998; Zera et al., 1993; Lentz et al., 1999; Maxwell et al., 2020). In the most common types of profile-analysis experiments, listeners are tasked with discriminating between a log-spaced complex tone with equal-amplitude components and a log-spaced complex tone with components that have equal amplitude except for the middle component, which has a higher amplitude. Here, we call the equal-amplitude components “background components” and the overall sound level of the background components the “background level”. We call the higher-amplitude component the “target component”, and the frequency of the target component the “target frequency”. In keeping with the literature (Green, 1988), we quantify the size of the amplitude difference between the target component and the background components in units of 20 X *log*_lO_(ΔA/A), which we refer to as “dB signal re: standard” (SRS), under the convention that the “signal” (*ΔA*) is the component added in-phase at the target frequency to increment the target component, and the “standard” (*A*) is the unincremented target component.

Performance in profile analysis is known to depend on a variety of stimulus parameters, including the total bandwidth of the tone complex (hereafter “bandwidth”), the number of components in the tone complex (hereafter “component count”), and the frequency spacing between components (hereafter “component spacing”). Holding bandwidth constant, performance is typically best when components are about ¼ octave apart and poorer when components are either further apart or closer together (Bernstein and Green, 1987; Lentz et al., 1999). Holding component spacing constant, increases in bandwidth, produced by adding additional flanking background components to the frequency edges of the stimulus, almost invariably result in improvements in performance (Green et al., 1983). Intriguingly, this effect of bandwidth is present even when the additional components fall well outside the critical band centered at the target frequency, which has been taken as evidence that listeners rely on information from “off-frequency” channels across a wide range of the tonotopic axis while performing the task.

Conventional interpretation of data from profile-analysis in roving-level paradigms suggests that two effects combine to result in the non-monotonic U-shaped relationship between component spacing and profile-analysis thresholds (Bernstein and Green, 1987). First, when components are distantly spaced, adding additional components initially allows the listener to make additional independent estimates of the background level of the stimulus, helping to mitigate the effects of level roving and leading to a gradual improvement in thresholds. Eventually, however, components become closely spaced enough that energetic masking effects begin to dominate and thresholds worsen again. These masking effects are usually interpreted under the power-spectrum model of masking (Patterson, 1976; Glasberg and Moore, 1990; Moore, 1995). However, several issues with this explanation of profile analysis raise questions about its validity. First, the physiological basis of the energy-detector system theorized by the power-spectrum model of masking is unclear; it is thought by some to be rooted in the excitation pattern, or average discharge rates across tonotopic peripheral responses (Sumner et al., 2018), implicitly perhaps in responses of non-saturating low-spontaneous-rate auditory-nerve (AN) fibers. However, many discrepancies between AN physiology and the power-spectrum model of masking have been noted (Carney, 2018). Consistent with this, prior work using a physiologically accurate computational model of the AN showed that decoding the responses of high-spontaneous-rate (HSR) AN fibers did not predict performance in a notched-noise masking task or a profile-analysis task (Maxwell et al., 2020). Second, it is unclear how the power-spectrum model of masking, which conventionally assumes that listeners use the output of a single auditory filter to perform the task, might be extended to handle the roving-level paradigm, which renders single-channel energy cues less reliable. Prior work tested the multichannel model of Durlach et al. (1986), but found that it could not account for data from different roving-level paradigms with a single set of parameters (Lentz et al., 1999).

Recently, an alternative theory of profile analysis has been advanced by Maxwell et al. (2020). In their theory, profile analysis is based on analysis of average rates at the output of a population of modulation-sensitive inferior colliculus (IC) neurons. Profile-analysis stimuli, due to peripheral interactions between nearby spectral components that are shaped by nonlinear auditory filtering and transduction, elicit slow fluctuations over time in the amplitude of AN responses. When the target component is incremented, these “neural fluctuations” are attenuated in channels tuned near the target frequency. This effect is due in part to the stimulus, as selectively amplifying the target component elicits lower modulation depth from interaction between the target component and nearby masker components, but the effect is also exaggerated by saturation of the inner-hair-cell input-output function (Carney, 2018). According to Maxwell et al. (2020), these changes in neural fluctuations due to the target increment could be decoded by modulation-sensitive IC cells to perform profile analysis. Multiple types of IC cells could be useful for decoding this cue (Kim et al., 2020); for example, cells with enhanced firing rates in response to modulation (band-enhanced, BE) would provide a target cue in the form of a decrease in discharge rate near the target CF, while cells with suppressed firing rates in response to modulation (band-suppressed, BS) would provide a target cue in the form of an increase in discharge rate near the target CF. The theory goes on to explain key trends in a profile-analysis task, including the resilience of human listeners to level roving, as the result of an interplay between cochlear tuning, nonlinear peripheral transduction, and modulation coding in the IC.

To help adjudicate between these competing explanations of profile analysis, we exploited the fact that changes to the frequency range of the stimulus have different predicted effects under the classical account and the neural-fluctuation account. In the former case, performance should remain fairly constant as a function of frequency, or even improve as a function of frequency, because auditory filters gradually sharpen at higher frequencies (Moore and Glasberg, 1987; Oxenham and Shera, 2003). In contrast, in the latter case, performance should degrade at sufficiently high frequencies because neural fluctuations elicited by profile-analysis stimuli will eventually be too fast to be encoded by fluctuation-sensitive auditory midbrain neurons (Kim et al., 2020). Limited prior evidence is consistent with the latter (Green et al., 1985), but more data is needed to provide a complete picture. We hypothesized that performance in profile analysis would be substantially worse for high target frequencies compared to low target frequencies, consistent with the neural-fluctuation theory.

To this end, we measured profile analysis as a function of target frequency, component spacing, and level-rove range. To improve our understanding of how processing of spectral profiles might be impacted by sensorineural hearing loss, these experiments included listeners with a wide range of sensorineural hearing loss. We also simulated neural responses to our stimuli at several stages of the auditory system to aid in interpretation of the psychophysical data. We found that performance degraded substantially for all listeners at high frequencies, consistent with the neural-fluctuation theory. Key trends observed in prior data (e.g., the non-monotonic U-shaped relationship between component spacing and profile-analysis thresholds) did not hold at high frequencies, raising doubts about classical interpretations of profile analysis. We also found that sensorineural hearing loss measurably worsened performance at 1 and 2 kHz and altered the relationship between thresholds and component spacing. Although we were unable to completely account for all trends in the data based on simulated neural responses, either at the level of the AN or at the level of the IC, models based on IC responses better captured key trends in the data with respect to target frequency, providing further evidence to support the idea that profile analysis is better interpreted as reflecting response patterns in the midbrain, rather than the auditory periphery.

## II. METHODS

### A. Participants and equipment

22 individuals participated in the experiment. All participants provided written informed consent prior to participation. All experimental procedures were approved by the University of Rochester Research Subjects Review Board. Participants received $15/hour as compensation for their time. Participants varied widely in age (21–77 years of age) and hearing status (−3–82.5 dB HL at the test frequencies). Hearing status was assessed by a standard audiogram (0.25–8 kHz), and participants were excluded if asymmetry exceeded 15 dB or if hearing loss was greater than 80 dB HL at any frequency. Individual listeners’ audiograms are shown in Figure 1.

**Figure 1:**
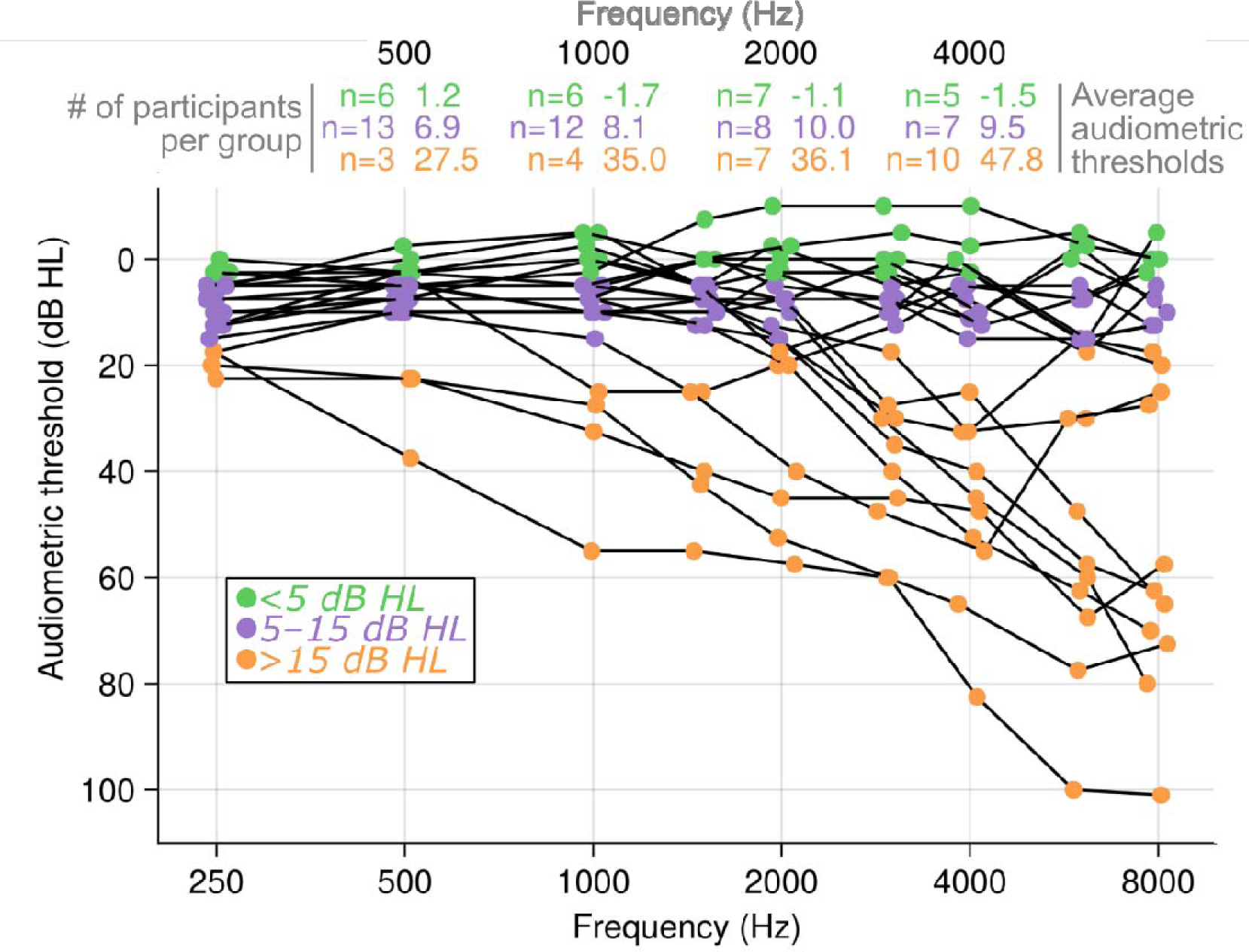
Listener audiograms. Audiometric thresholds for each listener as a function of frequency. Data corresponding to a single listener are connected by black lines. Thresholds at each test frequency are color coded to match the grouping variable used below when plotting behavioral data (see legend). Text above audiograms at frequencies of 0.5, 1, 2, and 4 kHz (the target frequencies used in the experiments reported below) indicates the number of listeners falling into each group and the average audiometric threshold of the group at the corresponding frequency.

During the experiment, the subjects were seated in a double-walled sound-attenuating booth. Listeners completed the experiment by interacting with a graphical user interface implemented with in-house code written in the M_ATLAB_ language (MathWorks, Natick, MA). Stimuli were presented to listeners diotically at a sampling rate of 48 kHz over TDH 39 earphones (Telephonics Corporation, Farmingdale, NY) via a MOTU M4 audio interface (MOTU Inc., Cambridge, MA) connected to a standard desktop computer. Test sessions were one or two hours in length.

### B. Behavioral procedure

Listeners heard three stimuli on each trial, separated by 300-ms intervals of silence. Each stimulus was an inharmonic complex tone with frequency components that were equidistant in log frequency. In the fixed-level paradigm, the stimuli were zero-phase. In the roved-level paradigm, phase offsets were independently selected for each component from a uniform distribution over [0, 2π]. The complex was centered on the target component and components spanned from 0.2× to 5× the target frequency (approximately a 4.6-octave range). The stimuli were 200-ms in duration, included 10-ms raised-cosine onset and offset ramps, and were presented at an overall sound level for the unincremented stimulus of 70 dB SPL on all trials (fixed-level condition) or over a range from 60–80 dB SPL selected randomly on each interval from a uniform distribution (roved-level condition). The first stimulus was always a reference stimulus (i.e., all components of the complex had equal amplitude). The second two were one reference stimulus and one target stimulus (i.e., middle [target] component had a higher amplitude), presented in random order. Listeners were tasked with indicating which of the latter two stimuli was the target.

In the first test session, listeners were introduced to the task with a suprathreshold stimulus, and a target level was identified at which the listener could perform the task at 100% correct. Listeners were then tested on a set of conditions for each target frequency, in random order across sessions. For each target frequency, five component-count conditions (5, 13, 21, 29, or 37 components) were tested. For each listener and target frequency, the tested increment sizes were configured by the experimenter in an attempt to sample the psychometric function efficiently and avoid spending significant time on the ends of the psychometric function. For each component count, listeners were tested in blocks of 20 trials at five different increment sizes. Listeners were tested in reverse order of increment size (i.e., a block of easy trials at a large increment size would be completed first, followed by a block of more difficult trials at a smaller increment size). Successive values of increment size were separated by 6 dB SRS. Each block began with two training trials; the responses to those trials were not recorded. If listeners answered the first five consecutive test trials correctly within a block, the block was terminated early, and testing moved on to a block for the next smaller increment size. If percent correct was less than or equal to 60% for two consecutive blocks, then testing for a given component count was stopped. Each block was repeated three times for each listener. Different component counts were tested in a randomized order.

For the fixed-level paradigm, which used a fixed level of 70 dB SPL in each interval, conditions included all combinations of component count (5, 13, 21, 29, or 37 components) and target frequency (0.5, 1, 2, or 4 kHz). For the roved-level paradigm, which used a random level for each interval uniformly selected from the range of 60–80 dB SPL, conditions included all component counts (5, 13, 21, 29, and 37 components) at a target frequency of 1 kHz. This testing yielded 20 conditions (4 target frequencies x 5 component counts) measured with the fixed-level paradigm in addition to another five conditions measured with the roved-level paradigm (5 component counts) for a total of 25 unique conditions.

It should be noted that stimuli were not amplified or altered in any way to compensate for hearing loss (i.e., all listeners heard the same stimuli with the same overall levels). Thus, at higher test frequencies, listeners who had substantial hearing loss were likely not able to hear the target component in a reference stimulus consisting of equal-amplitude components. Such listeners were likely still able to hear components on the low-frequency side of the stimulus, and the target component when incremented sufficiently, so they were sometimes still able to perform the task. Nevertheless, statistical analyses were constrained to only include data for a given participant and target frequency when the level of the target component (in a fixed-level reference stimulus consisting of equal-amplitude components) was above that participant’s audiometric threshold at the corresponding frequency. Data that were excluded from statistical analysis were also excluded from calculations of summary statistics shown below in figures. Figures showing individual data below do plot these individual excluded data, but they are marked with special symbols.

### C. Behavioral data analysis

Behavioral data were processed and analyzed in two stages. First, thresholds were estimated for each listener in each condition by pooling data across repeated measurements of the condition, computing the proportion of correct responses at each tested increment size, fitting a logistic psychometric function to the proportion-correct data using the Levenberg-Marquardt optimization algorithm, and then calculating the increment size that corresponded to 75.0% correct performance. Second, we fit a linear mixed-effects model using the R package *lme4* (Bates et al., 2015) to predict threshold values as a function of experimental parameters. In one model, we included fixed-level and roved-level 1-kHz data and used component count, size of rove range, and audiometric threshold at 1 kHz as fixed-effect variables. A full set of two- and three-way interactions between model terms was included. Component count and size of rove range were dummy coded as categorical variables, whereas audiometric thresholds were coded as continuous variables. Random-effect variables included random intercepts for each listener. In a second model, fixed-level data at all target frequencies were included but the roved-level data were excluded. For this model, component count, target frequency, and audiometric threshold at the target frequency were used as fixed-effects variables. Component count and target frequency were dummy coded as categorical variables, while audiometric threshold was used as a continuous variable. As before, random-effect variables included random intercepts for each listener. After fitting, standard evaluation plots (e.g., QQ plots and scale-location plots) were visually inspected to verify a suitable fit of the model and detect any issues during fitting. The fitted models were then analyzed with Type III ANOVAs (implemented in R via *car*; Fox and Weisberg, 2019) to assess omnibus significance of each predictor or interaction of predictors as well as a battery of contrast tests (implemented in R via *phia*; de Rosario-Martínez, 2015) to test *a priori* hypotheses as well as *post hoc* observations. In all cases, significance tests were corrected to control familywise error rates for each model separately using the Holm-Bonferroni correction (Abdi, 2010).

### D. Computational modeling

In order to better understand the behavior results and link them to known aspects of auditory neurophysiology, we simulated neural responses to our stimuli at various stages of the auditory system using phenomenological models of the AN and auditory midbrain (Nelson and Carney, 2004; Zilany et al., 2014; Carney et al., 2015) and analyzed those responses as described below.

#### 1. Auditory-nerve model

To simulate AN responses to our stimuli, we used the AN model of Zilany et al. (2014), a phenomenological model designed to replicate cat AN spike-train statistics in response to common laboratory sounds (e.g., pure tones, Gaussian noises), as well as arbitrary stimuli including speech sounds (Zilany and Bruce, 2006, 2007). The model captures several important features of peripheral physiology, including nonlinear cochlear filtering that is level- and CF-dependent, saturation and low-pass filtering in the inner hair cell (IHC), and adaptation and rate saturation associated with the synapse between the IHC and AN. The model was implemented using the original C source code with superficial modifications to allow for calls made from a custom codebase in Julia (Bezanson et al., 2017). Both the modified source code and the Julia wrapper are included in the code associated with this project (see below, [[link]]). Except when otherwise noted, the model was run at a sampling rate of 100 kHz, was configured to replicate human frequency-selectivity estimates (Shera and Oxenham, 2002; Ibrahim and Bruce, 2010), included fractional Gaussian noise in the IHC-AN synapse model, employed the approximate implementation of power-law adaptation in the IHC-AN synapse model to reduce overall compute time (Zilany et al., 2014), and had the C_IHC_ and C_OHC_ parameters set to values of 1. These simulations, and those described below, were conducted on Unix cluster computers administered by the University of Rochester Center of Integrated Research Computing.

#### 2. Auditory-midbrain model

The low-threshold HSR output of the Zilany et al. (2014) model was used as input to the same-frequency inhibition-excitation (SFIE) model to simulate neural responses from the cochlear nucleus (CN) and IC (Nelson and Carney, 2004; Carney et al., 2015). A single stage of the SFIE model accepts one excitatory input and one inhibitory input. The excitatory input and inhibitory input are instantaneous-firing-rate waveforms from the same CF, with the inhibitory input delayed by some amount (*d_i_*), hence the name “same-frequency”. The excitatory and inhibitory inputs are then convolved with different alpha functions of the form *P*(*t*) =*te^−t/r^*, a form inspired by the shape of ventral CN postsynaptic potentials (Nelson and Carney, 2004) that has the effect of temporally smoothing the inputs. The output of the model is then the half-wave-rectified weighted difference of the excitatory and inhibitory inputs. By calibrating the delay (*d_i_*), the excitatory and inhibitory time constants (*τ_e_*, *τ_i_*), and the excitatory and inhibitory weights (*w_e_*, *w_i_*), one can achieve a model that, for example, mimics rate tuning to amplitude-modulation frequency observed in IC neurons (Nelson and Carney, 2004). We simulated two different IC cell types, band-enhanced (BE) and band-suppressed (BS) neurons, with two slightly different configurations of the core SFIE building blocks. For BE simulations, a CN stage was first simulated by applying the SFIE algorithm to input HSR response waveforms with the parameters listed in the “CN” row of Table 1. Next, the IC stage was simulated by applying the SFIE algorithm again to the CN response waveforms with the parameters listed in the “BE” row of Table 1. BS neurons were simulated by first simulating BE neurons and then applying the SFIE algorithm but using the CN responses as the excitatory input and BE responses as the inhibitory input with the parameter values listed in the “BS” row of Table 1.

**Table 1:** Parameter values used in the SFIE simulations. Parameter values used for the SFIE model in the CN stage and in the two IC stages (BE and BS).

| | $\tau_e$ | $\tau_i$ | $d_i$ | $w_e$ | $w_i$ |
| --- | --- | --- | --- | --- | --- |
| **CN** | 0.5 ms | 2.0 ms | 1.0 ms | 1.5 | 0.6 |
| **BE** | 1.0 ms | 2.0 ms | 1.0 ms | 2.0 | 1.0 |
| **BS** | 1.0 ms | 2.0 ms | 1.0 ms | 0.2 | 20.0 |

#### 3. Normal-hearing simulations

The first step in the modeling procedure was to simulate population responses in four different model stages—AN-HSR, AN-low-spontaneous-rate (LSR), IC-BE, and IC-BS—in response to profile-analysis stimuli in every condition over a wide range of increment sizes. Populations were simulated for 91 CFs spanning logarithmically from ½ to 2 times the target frequency. The use of a log-symmetric frequency span and an odd number of CFs ensured that the middle CF was always exactly at the target frequency. In a given condition, responses to stimuli precisely matched to behavioral stimuli were independently simulated for each stage at increment sizes ranging from −45 to 5 dB SRS in 2.5 dB steps. Responses were simulated for 150 repeats of the reference stimulus and 150 repeats of the target stimulus at each of the increment-size steps and then averaged across time (including onsets) to yield an average discharge rate in each channel (hereafter a “response pattern”).

Second, each pair of reference and target response patterns (out of the 150 total at each increment size) was treated as a trial and decoded to arrive at a decision. It is worth noting that interstimulus-interval time was not simulated (i.e., simulated “trials” were consisted of reference and target response simulations conducted independently, rather than as a genuine pair with an intervening interstimulus interval). Two different decoding schemes were tested: single-channel, in which the interval evoking a higher discharge rate at the target CF (or a lower discharge rate, for the IC-BE model) was selected as the target, and template-based, in which the interval evoking a response pattern with a greater Mahalanobis distance from a template response pattern was selected as the target. The latter decoding strategy was motivated by Maxwell et al. (2020), who found that it was an effective strategy for decoding IC-BE responses in similar tasks. Conceptually, Mahalanobis distance is akin to a multi-dimensional d’ that accounts for the covariance of responses between CFs across trials. The template response pattern used for the template-based observer was generated by simulating responses to 500 reference stimuli and then estimating the average response pattern and the variance-covariance matrix. Separate templates were generated for fixed-level and roved-level conditions. Third and finally, psychometric functions were fit using the same procedure as for behavioral data to extract a threshold value at the 75.0% correct point for each tested model, observer, and stimulus condition.

#### 4. Hearing-impaired simulations

In addition to the main simulations described above, an additional set of simulations was conducted that factored in each participant’s unique hearing-loss configuration. The threshold-simulation process was repeated for the 1- and 2-kHz fixed-level conditions for each listener using a peripheral model customized to reflect their unique pattern of audiometric thresholds. Interpretable and significant effects of hearing loss were not seen for the 0.5- or 4-kHz target frequencies, so these target frequencies were not included in the hearing-impaired simulations. Specifically, the C_OHC_ and C_IHC_ model parameters, which respectively control the degree of simulated OHC and IHC dysfunction in the model, were adjusted for each CF to elevate model thresholds to match the listener’s audiometric thresholds. This was done using a variant of the *fitaudiogram2* function from the code release for the Zilany et al. (2014) auditory-nerve model (<u>link</u>) that was modified to use higher-resolution threshold data generated by linearly interpolating the original data along the log-frequency axis. In keeping with prior work (Zilany and Bruce, 2007; Bruce et al., 2013), it was assumed that up to 2/3 of audiometric losses could be attributed to OHC impairment and the remaining 1/3 to IHC impairment. To offset the additional computational cost associated with simulating results for 21 different listeners, these simulations were conducted at a reduced level of fidelity, and included only 61 CFs (rather than 91), 250 responses to estimate the template (instead of 500), and 50 simulated trials at each point on the psychometric function (rather than 150). Otherwise, simulations were conducted and analyzed exactly as for the normal-hearing simulations.

#### 5. Code availability

All code used to produce the analyses and figures in this paper is available at [[link]] under [[license]].

## III. RESULTS

### A. Behavioral results

Data from 1-kHz conditions are shown in Fig. 2. An ANOVA of thresholds across stimulus conditions and participants revealed significant main effects of component count [F(4, 166)=7.39, p<0.001] and level roving [F(1, 167)=136, p<0.001], as well as significant interactions between component count and level roving [F(4, 166)=9.04, p<0.001] and between component count and hearing loss [F(4, 166)=11.0, p<0.001]. No other effects achieved significance in the ANOVA (all p>0.118).

**Figure 2:**
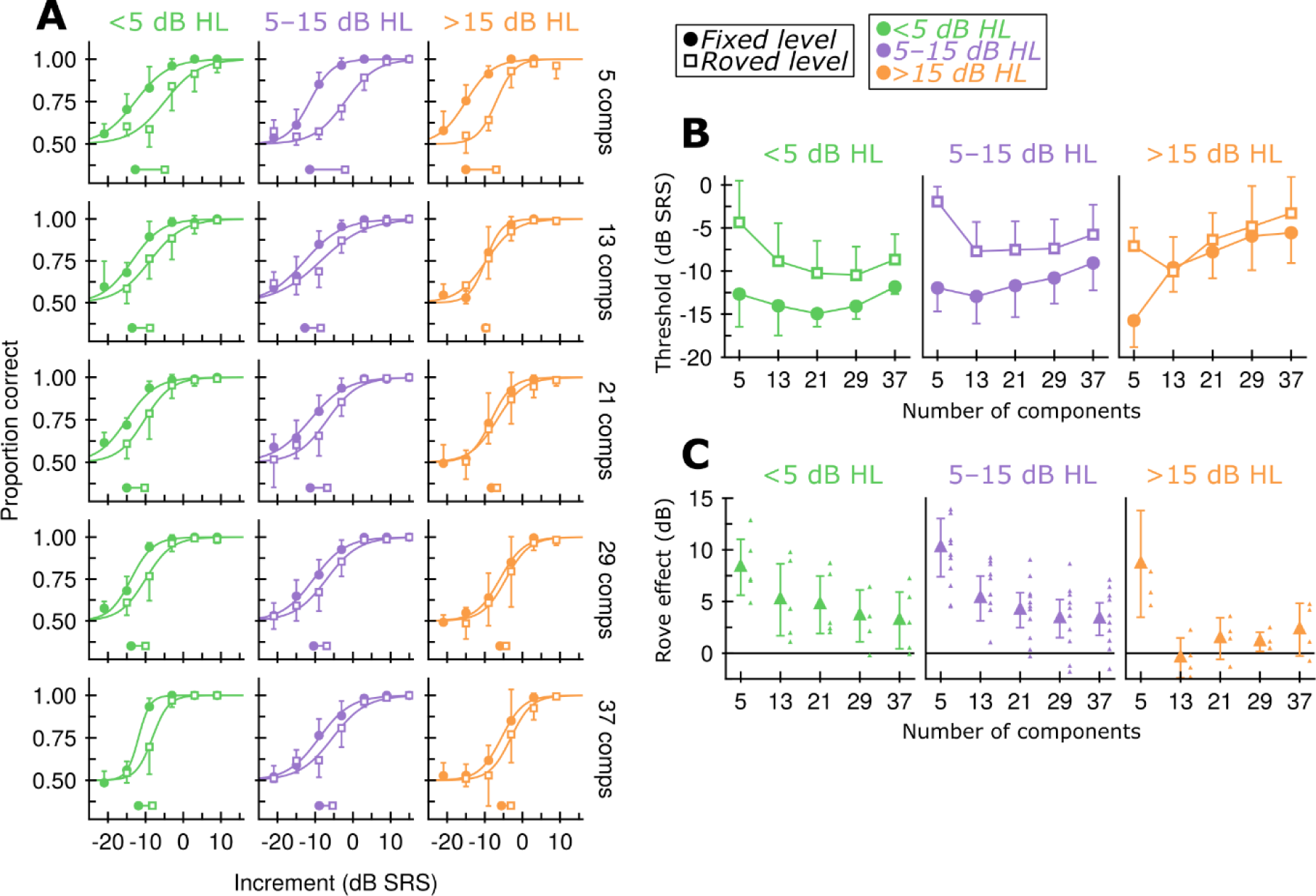
Effects of level roving and sensorineural hearing loss on profile-analysis. **A.** Group-average psychometric functions for listeners for the 1-kHz target-frequency condition. Listeners were grouped based on hearing loss at 1 kHz: less than 5 dB HL (green), 5–15 dB HL (purple), or greater than 15 dB HL (orange). Different columns show different groups of listeners, while different rows show different stimulus component counts. Fixed-level data are shown as filled circles, while roved-level data are shown as unfilled squares. Error bars indicate ±1.96 standard error of the mean. Markers below the psychometric functions indicate thresholds at the 75%-correct point of the psychometric function. **B.** Group-average thresholds as a function of the number of stimulus components. Color and marker type indicate the same as in A. Error bars indicate ±1.96 standard error of the mean. **C.** Group average rove effect (difference between roved-level and fixed-level thresholds) as a function of the number of stimulus components. Color indicates the same as in A. Error bars indicate ±1.96 standard error of the mean. Small markers indicate rove-effect magnitudes for individual participants.

We will first turn our attention to the effects of level roving on profile analysis. Averaged across all conditions and listeners, roved-level thresholds were significantly higher than fixed-level thresholds [estimated difference = 4.66 dB, F(1, 167)=137, p<0.001]. The effect of level roving varied significantly as a function of component count. This interaction can be seen readily in Fig. 2b, which compares average thresholds for the fixed-level and roved-level conditions as a function of component count, and Fig. 2c, which shows the change in threshold produced by level roving as a function of component count. The effect of the rove was significant and large for the 5-component stimulus [estimated rove effect = 9.32 dB, F(1, 166)=111, p<0.001], while for other component counts the effect of level roving was significant [estimated rove effect = 3.07–4.13 dB, all p <0.001] but smaller than for the 5-component stimulus [estimated difference of rove effect = 5.19–6.25 dB, all p<0.001].

Next, we will turn our attention to the effects of sensorineural hearing loss. We found that, averaged across fixed- and roved-level data, the effect of hearing loss (quantified as the audiometric threshold at 1 kHz) was significant for 21-component stimuli [F(1, 27.9)=4.17, p=0.051], 29-component stimuli [F(1, 27.9)=7.70, p=0.010], and 37-component stimuli [F(1, 27.9)=4.58, p=0.041], but not for 5- or 13-component stimuli (all p>0.263); effects of hearing loss for the 21-, 29-, and 37-component stimuli were significantly larger than for the 13-component stimuli (all p < 0.029). In other words, audiometric thresholds were significantly related to profile-analysis thresholds for spectrally dense stimuli, but not for spectrally sparse stimuli. Consistent with a significant interaction between hearing loss and component count, the overall shape of the threshold vs. component-count curves changed with increasing hearing loss. For listeners with less than 15 dB HL at 1 kHz, thresholds were fairly flat or exhibited a non-monotonic U-shape as a function of component count. In contrast, for listeners with greater than 15 dB HL, thresholds were monotonically increasing with component count. Although the interaction between level roving and hearing loss was not significant in our ANOVA, visual inspection of Figs. 2b and 2c suggests that listeners with audiometric thresholds higher than 15 dB HL at 1 kHz exhibited *less* sensitivity to the level rove than other listeners. This trend was unexpected, given prior work showing that listeners with sensorineural hearing loss are *more* sensitive to level roving in a different task (Leong et al., 2020). The trend appeared to be driven by elevated fixed-level thresholds for hearing-impaired listeners (except for the 5-component stimulus).

Data from fixed-level conditions across all target frequencies are shown in Fig. 3 (the fixed-level data for 1 kHz shown in Fig. 3 are the same as the fixed-level data shown in Fig. 2). The ANOVA of this dataset revealed significant main effects of component count [F(4, 360)=15.9, p<0.001], frequency [F(3, 365)=115, p<0.001], and hearing loss [F(1, 376)=13.0, p<0.001]. Significant two-way interactions were detected between component count and frequency [F(12, 360)=2.18, p=0.012], component count and hearing loss [F(4, 360)=6.41, p<0.001], and frequency and hearing loss [F(3, 370)=10.7, p<0.001]. A significant three-way interaction between all model terms (component count, target frequency, and hearing loss) was also found [F(12, 360)=2.87, p<0.001].

**Figure 3:**
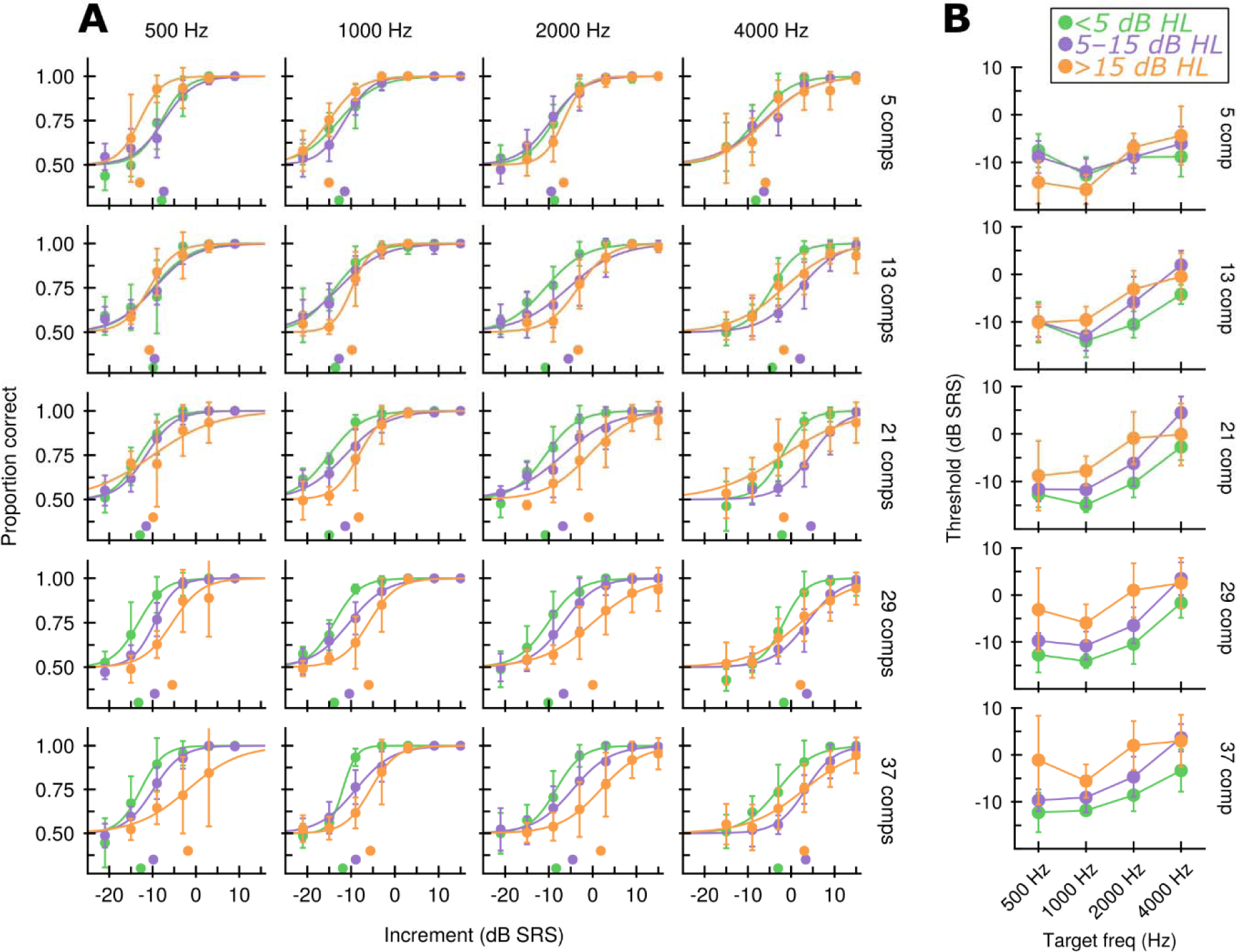
Effect of target frequency on profile-analysis. **A.** Group-average psychometric functions from the fixed-level conditions for listeners with less than 5 dB HL (green), 5–15 dB HL kHz (purple), or greater than 15 dB HL (orange) at the target frequency. Different columns show target frequencies, while different rows show different stimulus component counts. Error bars indicate ±1.96 standard error of the mean. Markers below the psychometric functions indicate thresholds at the 75%-correct point of the psychometric function. **B.** Group-average thresholds as a function of target frequency. Color indicates the same as in A. Error bars indicate ±1.96 standard error of the mean.

We will first focus our attention on the effects of target frequency on profile analysis. The relevant data are shown in Fig. 3b. On average, thresholds were not significantly different between 0.5 and 1 kHz [estimated difference = 0.790 dB, F(1, 366)=1.76, p=1.00], but significantly worse thresholds were observed at 2 kHz compared to 1 kHz [estimated difference = 5.64 dB, F(1, 366)=119, p<0.001] and at 4 kHz compared to 2 kHz [estimated difference = 4.94 dB, F(1, 366)=87.4, p<0.001]. This consistent worsening of thresholds beyond 1 kHz is consistent with limited prior data on the effects of target frequency on profile analysis (Green and Mason, 1985). Consistent with a significant interaction between component count and target frequency, the elevation of thresholds at higher frequencies was apparent for most component counts but less clear for the 5-component stimulus.

Next, we turn to effects of hearing loss, and interactions between hearing loss and target frequency, which are visualized in Fig. 4. Averaging across component count, we found that audiometric thresholds and profile-analysis thresholds were significantly correlated at 2 kHz [F(1, 375)=17.2, p=0.001], but not at other frequencies (all F < 4.09, p > 0.789). This correlation was significantly larger at 2 kHz than at other target frequencies (all F > 10.59, p < 0.002). We observed a significant effect of hearing loss for the combined fixed- and roved-level 1-kHz data above (Fig. 2), but not for the fixed-level 1-kHz data alone; however, the present analysis has less overall statistical power when considering each frequency condition separately, and there was still an overall positive trend between audiometric thresholds at 1 kHz and fixed-level profile-analysis thresholds at 1 kHz, despite not reaching significance. The significant interaction between target frequency and hearing loss likely reflects several contributing factors. At low target frequencies (0.5 and 1 kHz), our listeners had (relatively) little hearing loss. For example, at 1 kHz, the median audiometric threshold was 7.5 dB HL, and 75% of participants had less than 10 dB of loss. This small range of audiometric thresholds limited the extent to which we could measure a correlation between hearing loss and profile-analysis thresholds. At 4 kHz, audiometric thresholds spanned a much larger range, but the task was quite difficult overall (even for normal-hearing listeners). Moreover, some hearing-impaired listeners exhibited unexpectedly good performance at 4 kHz, possibly due to issues with the audibility of the stimulus for some listeners (see Discussion), which disrupted a linear relationship between audiometric thresholds and profile-analysis thresholds at 4 kHz. Collectively, these factors combined to make 2 kHz the best target frequency to observe an effect of hearing loss on profile analysis thresholds. A recent study on format-frequency discrimination that included many of the same listeners likewise observed significant relationships between audiometric thresholds at 2 kHz and formant-frequency discrimination at 2 kHz, but not between audiometric thresholds at 0.5 kHz and formant-frequency discrimination at 0.5 kHz, possibly for similar reasons (Carney et al., 2023).

**Figure 4:**
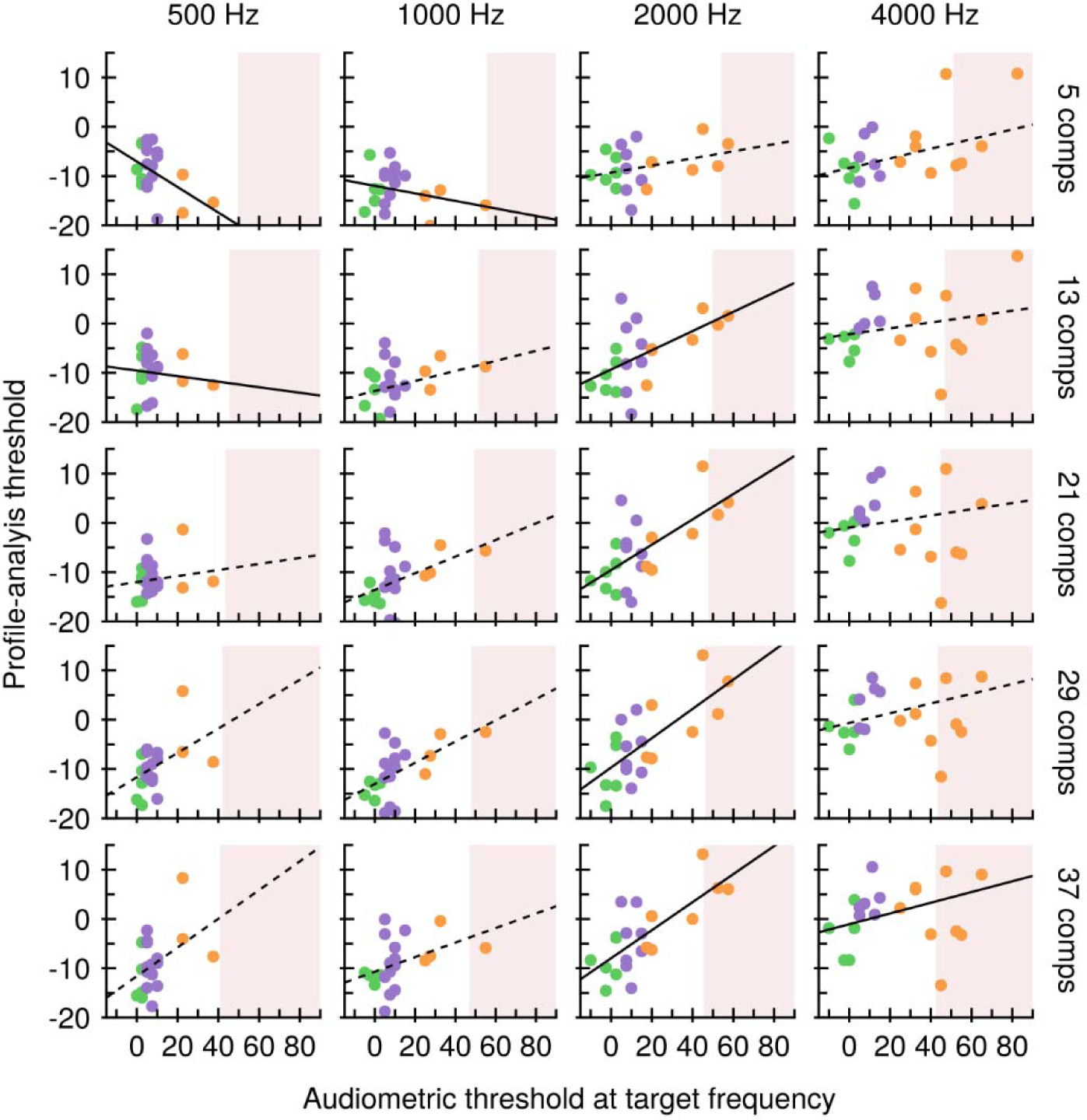
Correlations between audiometric thresholds and profile-analysis thresholds. Individual profile-analysis thresholds as a function of audiometric threshold at the target frequency. Different columns indicate different target frequencies, while different rows indicate different numbers of stimulus components. Color indicates which group points were assigned to in Figs. 1 and 2 based on audiometric threshold. Black lines indicate linear regressions through the data, with dashed lines indicating relationships that are not significant and solid lines indicating relationships that are significant. The red region indicates audiometric thresholds for which the target-component sound level was nominally below absolute threshold.

Hearing loss also interacted significantly with component count, as in the model analyzing the 1-kHz data above. Specifically, we found that the effect of hearing loss was significant on average for 37-component stimuli [F(1, 378)=5.46, p=0.020] and 29-component stimuli [F(1, 378)=6.27, p=0.0127]. For the 21- and 13-component stimuli, no significant effect of hearing loss was detected (all F < 1.011, p > 0.315). Intriguingly, for the 5-component stimuli, a significant effect of hearing loss was detected [F(1, 378)=15.4, p<0.001], but the association was *negative*. That is, listeners with greater degrees of hearing loss had *better* thresholds on average for the 5-component stimuli. This unexpected trend was driven by results in the low-frequency conditions, where the effect of hearing loss in the 5-component condition was significant [0.5 kHz, F(1, 375)=18.7, p<0.001; 1 kHz, F(1, 375)=10.5, p=0.001], as opposed to the high-frequency conditions, where no such significant effect was detected (all F < 1.99, p > 0.160).

### B. Computational results

#### 1. Normal-hearing computational results

In order to better understand what aspects of auditory processing might give rise to the patterns observed in our behavioral data, we simulated responses to our behavioral stimuli using a phenomenological model of the auditory periphery and subcortex (Nelson and Carney, 2004; Zilany et al., 2014; Carney et al., 2015) and decoded them to predict behavioral thresholds. Figure 5a depicts the signal flow of our computational model. Figure 5b shows responses to standard auditory characterization stimuli (e.g., pure tones, modulated noises) for each model. As can readily be seen, the HSR, BE, and BS models all share similar low absolute thresholds, limited dynamic ranges, and saturating rate-level functions, whereas the LSR model has a higher absolute threshold, a wider dynamic range, and a sloping rate-level function (Fig. 5b, top row). The dashed gray line in each subpanel indicates the per-component level used for the 21-component stimulus in our experiments, and thus serves as an indicator of whether or not we expect HSR/LSR rates to be saturated when responding near stimulus components. All of the models have sharp frequency tuning at low sound levels (Fig. 5b, middle row), but only the BE and BS models show clear tuning to modulation frequency (Fig. 5b, bottom row). The modulation frequency that elicited the highest rate for the BE neuron, or its best modulation frequency (BMF), was 74 Hz, comparable to the population BMF in rabbit (Kim et al., 2020).

**Figure 5:**
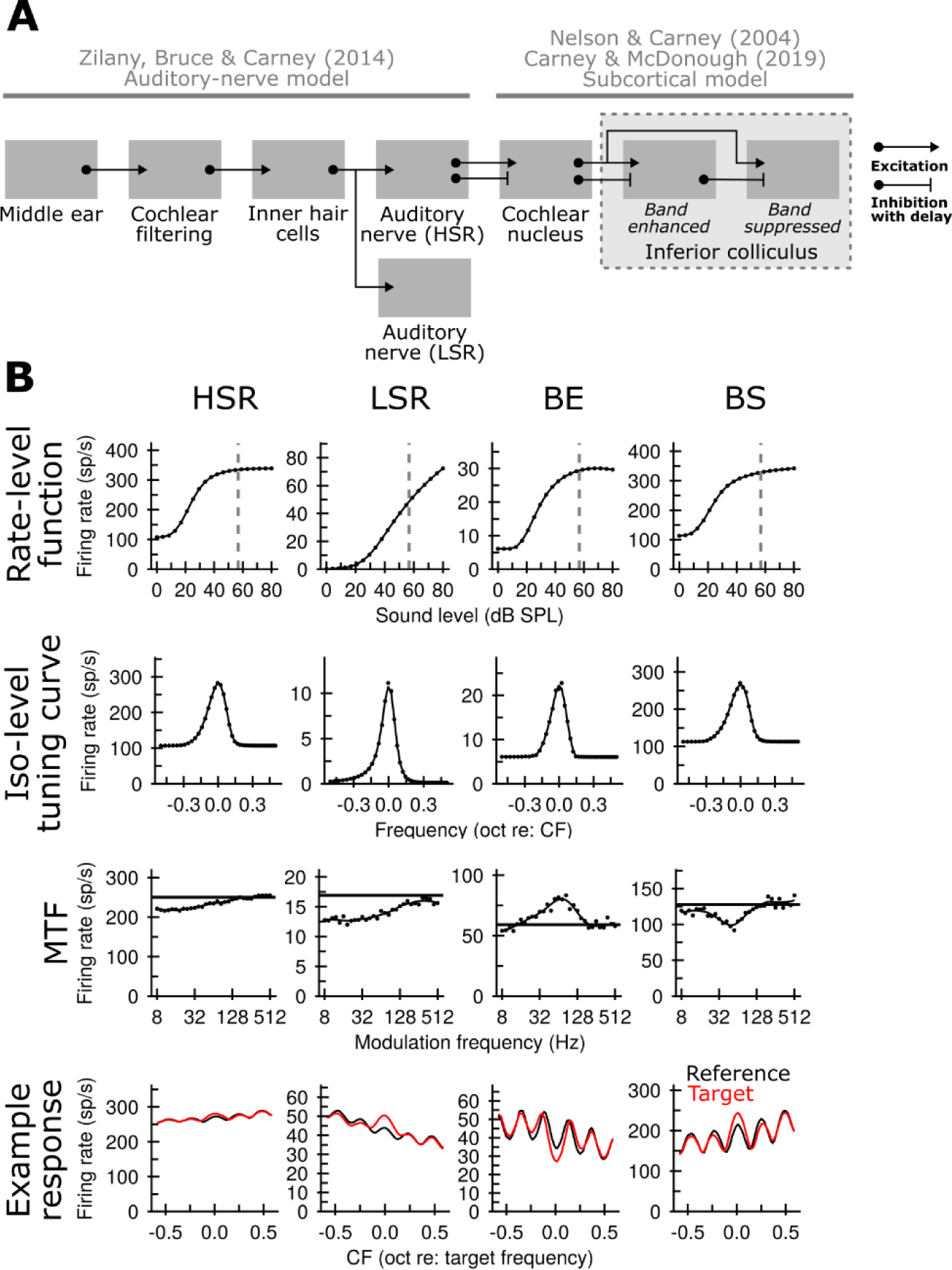
Modeling pipeline and key features of modelled stages. A. Schematic depicting signal flow in the auditory computational model used to conduct simulations. B. Key model responses properties and examples profile-analysis responses for model units with CF fixed at 2 kHz. Top row: rate-level functions showing average rates in response to a 100-ms pure tones at over a range of levels. Second row: iso-leve tuning curves showing average rates in response to a 30 dB SPL pure tone at a range of frequencies near CF. Third row: modulation transfer functions (MTFs) showing average rate in response to 500-ms Gaussian sinusoidally amplitude-modulated (SAM) noises with a spectrum level of 20 dB SPL and 100% modulation depth over a range of modulation frequencies. Bottom row: example responses to a profile-analysis reference tone (solid) and target tone (dashed) at an increment size of –10 dB SRS for a target frequency of 2 kHz (color online).

The bottom row of Fig. 5b shows example responses to the profile-analysis stimulus in each model. In response to a moderate increment size of –10 dB SRS (a near-threshold value for many listeners and conditions), the HSR response pattern was fairly flat across model CFs, clear peaks were visible in the LSR and BS responses, and a clear trough was visible in the BE response. The results for the HSR and LSR models are easy to understand. For the HSR model, response patterns are flat, and similar for the reference and target stimuli, because the HSR model rate is limited by rate saturation; that is, the HSR model cannot code the increment via rate, because its average rate is already saturated in response to the reference stimulus. The LSR model, in contrast, does not have a saturated average rate at the levels used, and therefore has a clear peak in the response pattern for CFs tuned near the target tone. Results for the BE and BS models are somewhat more complex, because these models are sensitive to both energy and temporal features, such as the neural fluctuations on their inputs. In response to the reference stimulus, the CN inputs (driven by HSR responses) to both the BE and the BS model contain fluctuations that arise from beating stimulus components and are shaped by peripheral filtering and transduction. When the target component is incremented in level, responses in the AN undergo capture and have weaker fluctuations. BE neurons, which are enhanced by modulation, show a commensurate decrease in rate. BS neurons, which are suppressed by modulation, show a commensurate increase in rate.

Modeling results are shown in Figs. 6-8. Figure 6 shows the changes in response patterns associated with a −5 dB SRS increment for each simulated model in all the stimulus conditions. Several qualitative observations are worth highlighting before considering quantitative analyses below. First, it is clear from Fig. 6A that little task-related information was contained in HSR fiber average discharge rates. Except in off-target-CF channels tuned between distant stimulus components, where effective input levels to the AN were low after cochlear filtering (i.e., 5-component stimulus, top row), rate saturation of the HSR fibers (Fig. 5B) virtually eliminated the ability to code the level increment in terms of average discharge rates. In the following figures, HSR results are thus excluded for brevity, though it is worth noting that the instantaneous discharge rates of HSR AN fibers are the inputs to the IC models.

**Figure 6:**
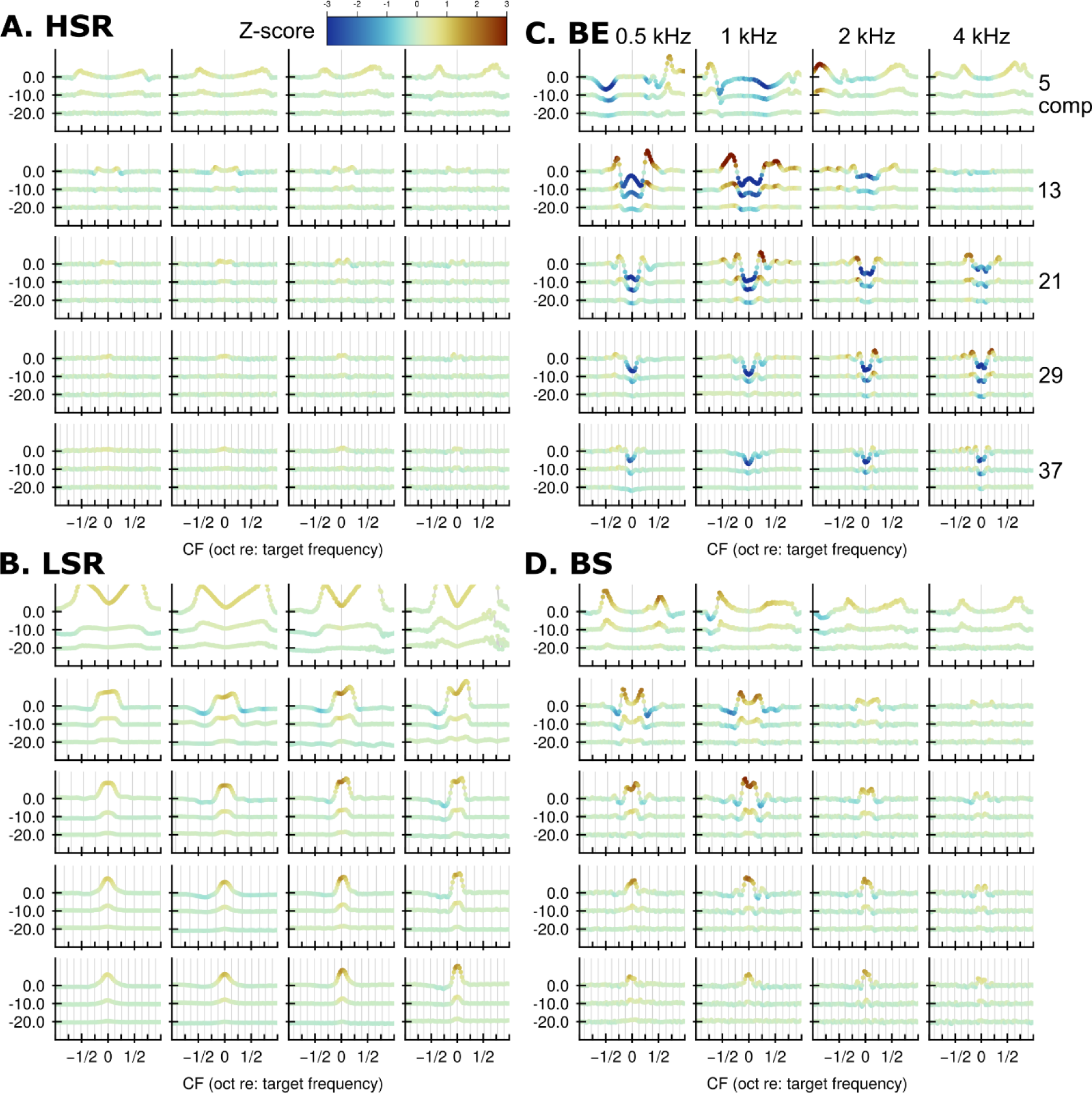
Example response profiles. A. Average differences between target and reference response patterns in each condition for the HSR model at various increment sizes. Results for different increment sizes are placed at different vertical offsets. Deviation from the vertical offset at each CF indicates the average change in rate in response to the increment, expressed as a percentage relative to the average reference rate. Each vertical offset corresponds to 50% change in rate. Color at each CF indicates the mean percent difference divided by its standard deviation, thus providing a measure of how consistent the changes in rate were from trial to trial. Simulations with level roving are shown; simulations without level roving resulted in nearly identical mean curves and are not shown. B. Same as A, but for the LSR model. C. Same as A, but for the BE model. D. Same as A, but for the BS model.

**Figure 7:**
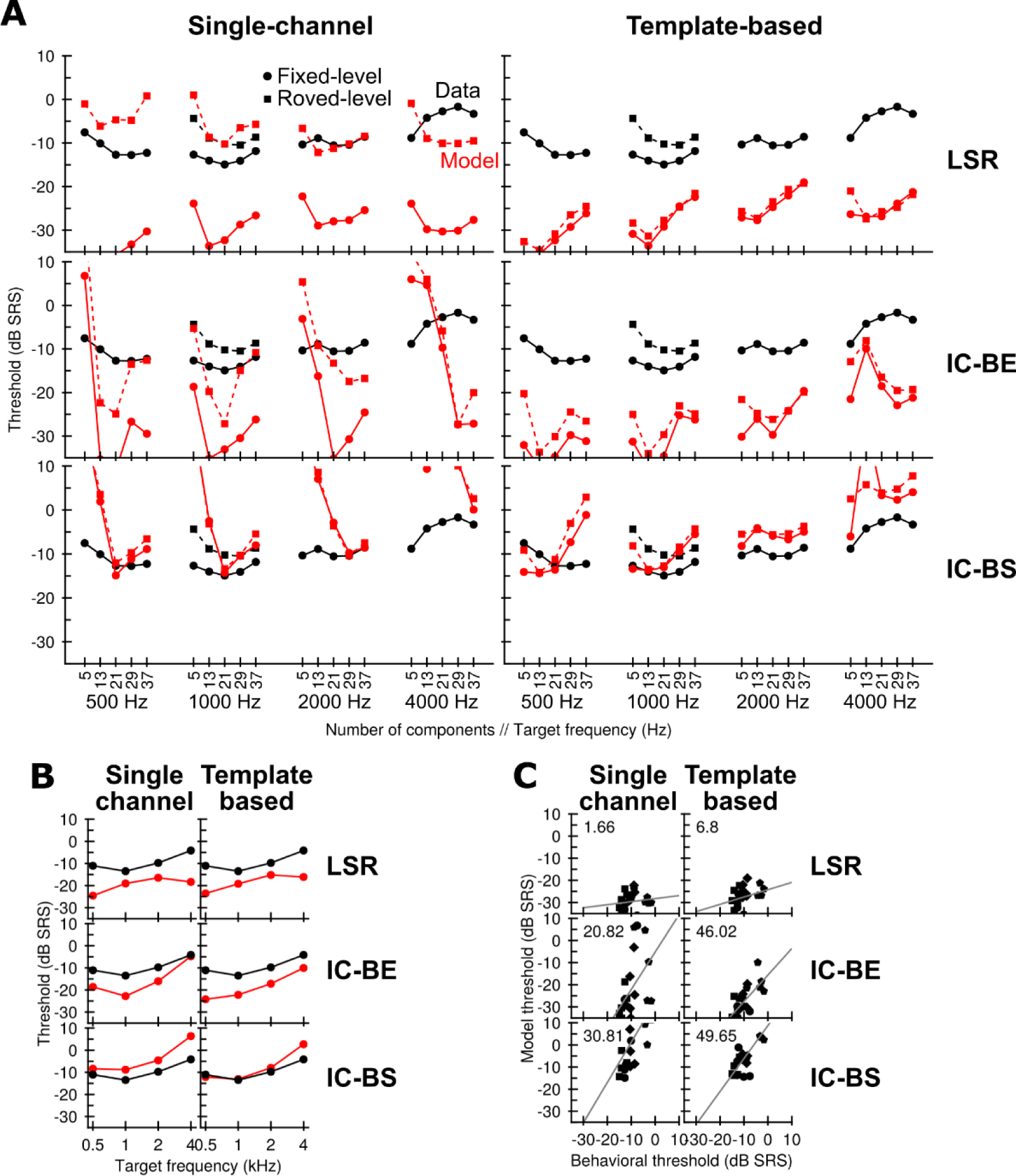
Predicted thresholds vs behavioral thresholds for listeners with less than 5 dB HL at the target frequency. A. Mean behavioral fixed-level thresholds (black circles) and roved-level thresholds (black squares) in each condition (x-axis) versus modeled fixed-level thresholds (red circles) and roved-level thresholds (red circles) under different observers (columns) and auditory-model stages (rows). B. Thresholds for fixed-level behavior (black) and models (red) averaged across all component counts to emphasize overall trends with respect to target frequency. Columns and rows are laid out in the same way as in A. C. Correlations between fixed-level behavioral thresholds (x-axis) and modeled thresholds (y-axis). Marker shape indicates target frequency. The gray line marks a linear model fit to predict model thresholds as a function of behavioral thresholds, while the number in the upper left corner of each panel indicates the percent variance explained by said linear model.

**Figure 8:**
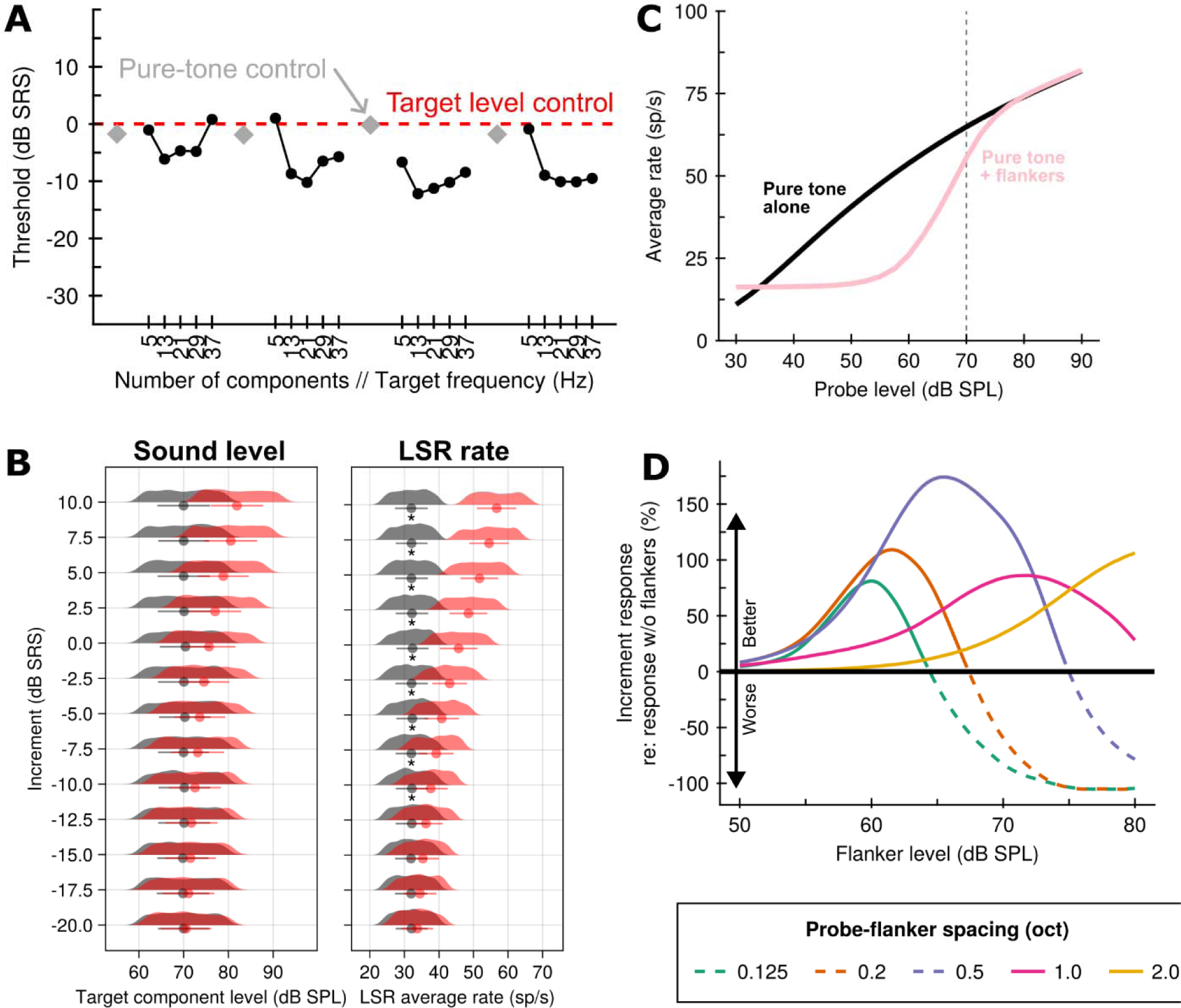
Measurements of LSR level-increment responses. A. Modeled roved-level single-channel LSR thresholds in each condition (black) as compared to predicted thresholds based on selecting the interval with the higher target level on each trial (red dashed line) or modeled single-channel LSR thresholds for a roved-level profile-analysis stimulus with no masker components (i.e., pure tone). B. Left, density kernel histograms of target-component sound levels for reference stimuli (gray) or target stimuli (red); right, density kernel histograms of average LSR discharge rates in the channel tuned nearest to the target frequency for reference stimuli (gray) or target stimuli (red). The stimulus used here was a 21-component 2-kHz stimulus. C. Rate-level functions for the LSR model fiber at a CF of 2 kHz measured for a pure tone in isolation (black) or a pure tone with two flankers, each at a level of 70 dB SPL, ½ octaves above and below the probe frequency (pink). D. Size of the “increment response”, or change in rate associated with a 0 dB SRS increase in the sound level of the probe tone, as a function of the sound level of two flanking pure tones positioned above and below the probe tone at different spacings (color). The size of the increment response is quantified in terms of percentage change relative to the increment response with no flanking tones.

LSR fibers, shown in Fig. 6B, did not suffer from this same limitation and had response patterns with robust peaks near target CF in most conditions. Peaks in LSR response patterns were largest for 13/21 components and were somewhat smaller for 29/37 components, consistent with the *a priori* expectation that more components entering into the cochlear filter tuned near the target frequency should result in a smaller increment response. Peaks in LSR response patterns were fairly stable over target frequency, although the 4-kHz target frequency generally elicited the largest response, consistent with sharper cochlear filters in the model at higher frequencies (Zilany et al., 2009; Zilany et al., 2014). It is evident from the span of the lines denoting the range of ±1 standard deviation around the mean that the LSR average discharge rates were highly variable under the level rove. This variability would limit how well the LSR rates could be used to code the increment; quantitative analysis of the consequences of this variability are presented below.

Figures 6C and 6D show that BE and BS response profiles, respectively, reflected the increment size in very different ways than the LSR response profiles. BE response profiles generally showed a dip in rates of neurons with CFs near the target frequency, whereas BS response profiles generally showed a peak for CFs near the target CF. For both BE and BS patterns, the precise shape of the dip or peak near target CF depended in complicated ways on target frequency and component count. Generally speaking, dips/peaks were most prominent for intermediate component counts and for lower target frequencies. At the 4-kHz target frequency, dips/peaks were quite small or altogether nonexistent. This degradation at high target frequencies was a consequence of increases in modulation rates elicited by beating between stimulus components near the higher target frequencies. At sufficiently high modulation frequencies, elicited neural fluctuations fell outside the modulation-coding range of the IC model (a limitation that is consistent with known IC physiology; Kim et al., 2020). In comparison to the LSR response patterns, the BE response patterns in particular showed substantially more stability with respect to the randomly varying sound level.

Figure 7 shows modeled thresholds versus behavioral thresholds for participants with less than 5 dB HL at the target frequency across all conditions and tested combinations of observer strategies and auditory-model stages. We will first evaluate the models in terms of their ability to account for trends with respect to target frequency and component count, beginning with the results of the LSR model. Decoding LSR response patterns led to different predictions with respect to target frequency than were seen in behavioral data. Both the single-channel LSR model and the template-based LSR model erroneously predicted a worsening in performance from 0.5 to 1 kHz target frequency and an improvement in performance from 2 to 4 kHz target frequency, but in behavior the opposite trends were observed (Fig. 7B). As mentioned previously, this moderate improvement at higher frequencies is consistent with sharper relative cochlear tuning at higher CFs in the model improving the fidelity of rate coding at higher target frequencies. The LSR model also failed to predict the significant interaction between component count and target frequency seen in the behavioral data; whereas behavioral data showed flat or U-shaped curves as a function of component count at lower target frequencies and an inverted-U shape at 4 kHz, the LSR models maintained a consistent U-shaped curve at all target frequencies.

The single-channel BE and BS models were too sensitive to changes in component count as compared to human listeners, but template-based decoding of the BE and BS response patterns led to thresholds that were more uniform across component count. This model result stands in contrast to the LSR results, which showed fairly similar shapes for single- and multi-channel decoding, suggesting that information contained in “off-target-CF” responses contributed more to decoding BE and BS responses than LSR responses. This difference likely arose in part because, unlike for the LSR model (barring the 5-component condition), the channel tuned to the target frequency (i.e., CF = target frequency) was not always the most informative for the BE and BS models (Fig. 6). Template-based thresholds for the BE model did not show the same trends as listeners with respect to component count, while template-based thresholds for the BS model roughly resembled behavioral data at 2- and 4-kHz target frequencies. However, at 0.5- and 1-kHz target frequencies, the BS model predicted degradation in thresholds with increasing component count that exceeded what was observed in behavior. Importantly, although neither the BE or the BS model accurately predicted the effects of component count on thresholds, both models showed trends with target frequency that were similar to behavioral data (Fig. 6b). In particular, decoding BE and BS response patterns by either tested method resulted in thresholds that were most elevated at 4 kHz, consistent with behavioral data.

Next, we turn our attention to considering the effects of level roving. One of the most curious features of profile analysis is that thresholds are relatively robust to a substantial level rove (Fig. 2). The mechanisms underlying this resilience are unknown, but the present behavioral and modeling results highlight several possibilities. One possibility is that rove resistance could be provided by information from off-target-CF responses that helps discount the confounding effects of level variation. Single-channel LSR responses were rendered less reliable by level roving for the same reason that single-channel LSR rates can code the target increment in the fixed-level condition: variations in level lead to variations in LSR rate. Commensurate with this, single-channel LSR thresholds were elevated substantially by level roving (sometimes by as much as 30 dB or more; Fig. 6A, left column, red squares vs. circles). However, the template-based LSR thresholds were unaffected by level roving (Fig. 6A, right column, red squares vs. circles). This large difference was because variations in overall level led to changes in rate that were correlated across the neural population (i.e., an increase in overall level, generally speaking, led to increased LSR rates in most channels), whereas variations in increment size led to changes in rate that were localized to the target CF, and the Mahalanobis-distance metric that underlies the template-based observer was able to distinguish these two types of rate changes. Practically, this result suggests that the LSR fibers contained enough information to achieve resistance to a moderate level rove, but this resistance required appropriate consideration of responses from multiple channels (i.e., it required a suitable population-decoding scheme).

However, the results of the BE and BS models suggest another intriguing possibility. In many cases, single-channel BE and BS thresholds were unaffected by roving, or at least single-channel roved thresholds were quite good and often better than behavioral data. BE and BS models are sensitive to sound level, but, like the HSR fibers that drive them and from which they inherit their rate-level functions for pure tones, changes in sound level only result in changes in their rates at low sound levels (Fig. 5B). At higher levels, such as those used in the present experiments, simple increases in energy do not further increase rate. This rate saturation produces a sort of resistance to roving, insofar as it makes rates more stable with respect to variation in sound level. For a neuron exclusively sensitive to energy, this rove resistance would be counterbalanced by poor coding of the increment, and as such would be of little practical benefit. In contrast, coding the target increment remained possible for the BE and BS models because changes in neural fluctuations on their inputs still resulted in changes in discharge rate, even when the models had saturated rates as quantified by a pure-tone rate-level function. This result suggests that rate saturation, like other auditory nonlinearities, may be a “feature”, rather than a “bug”, of the auditory system (Carney, 2018), helping to make rates more stable with respect to irrelevant features (in this case, overall sound level) so that relevant features (in this case, relative target level) can be reliably coded, even when irrelevant features vary unpredictably.

A third factor to consider is revealed by the fact that roved LSR single-channel thresholds fell within the range of behavioral thresholds and exceeded expected performance based on an energy-detector model (Fig. 8A). Follow-up simulations strongly suggested that cochlear suppression, which is included in the AN model (Zilany et al., 2014), gave rise to this effect. When simulations were repeated with a 1-component stimulus (i.e., pure-tone level discrimination with a level rove), model thresholds closely matched the theoretical performance of an energy-detector model that was limited only by the presence of a level rove (e.g., Green, 1988, Appendix A). Figure 8B compares the distributions of target component level to distributions of LSR average rates at the target CF for reference and target stimuli with 21 components. As can be seen, LSR target and reference rates were more separable at smaller increment sizes than were the target component levels themselves; in other words, LSR target and reference rates were more different than would be expected based on differences in stimulus energy at the target frequency. Rate-level functions were then simulated for a pure tone in isolation and for a pure tone with flanking tones similar to profile-analysis masker components. These simulations revealed that, due to cochlear suppression, the LSR model’s rate-level function was steeper near levels used in the experiment when flankers were present than when flankers were absent (Fig. 8C). The suprathreshold enhancement of a fixed 0-dB SRS increment response was maximized when the flankers were comparable in level to the probe and approximately ½ octave away from the target (Fig. 8D). In brief, peripheral nonlinearities in the AN model enhanced coding of relative level increments in LSR fibers (and presumably also in other stages of the model).

#### 2. Hearing-impaired computational results

In Fig. 7, we compared results from a normal-hearing computational model to results from listeners with audiometric thresholds better than 5 dB HL at the target frequency. Next, we consider whether our modeling framework can account for our behavioral data across the full range of sensorineural hearing loss among our participants. Recent work has shown that adjusting peripheral models to account for individual variability in audiometric thresholds can improve predictive accuracy (Zaar and Carney, 2022; Carney et al., 2023). Accordingly, we created an individualized peripheral model for each participant and then repeated a subset of our simulations with the models (1- and 2-kHz target frequencies with a fixed-level, where significant effects of hearing loss were observed, but thresholds were measurable; Fig. 3). This customization was achieved by altering the C_OHC_ and C_IHC_ parameters in each channel of the peripheral model to account for the participant’s audiometric thresholds (see Methods for details).

Figure 9A shows correlations between behavioral thresholds and model thresholds with individualized audiograms for the 2-kHz fixed-level condition, pooled across different component counts. For subjects with less than 15 dB of hearing loss at the target frequency (i.e., groups “< 5 dB HL” and “5-15 dB HL”), adding individualized audiograms did not predict the individual variability in the dataset (< 5.1% variance explained by any model/observer combination). In contrast, when considering only listeners with more than 15 dB of hearing loss at the target frequency, individualized audiograms explained substantial variability in the dataset, with the BS-profile-based model able to account for the largest portion of variance (61.0%), the LSR-profile-based and single-channel models both able to account for slightly less (51.4% and 51.7%, respectively), and the BE-profile-based model able to account for the least (34.1%; other decoding schemes applied to the BE model outputs accounted for little variance). Although considerable variance remained unexplained, these proportions of variance explained are comparable to similarly derived values reported in Carney et al. (2023) and suggest that the present modeling framework is a useful way to investigate the effects of sensorineural hearing loss at the individual level.

**Figure 9:**
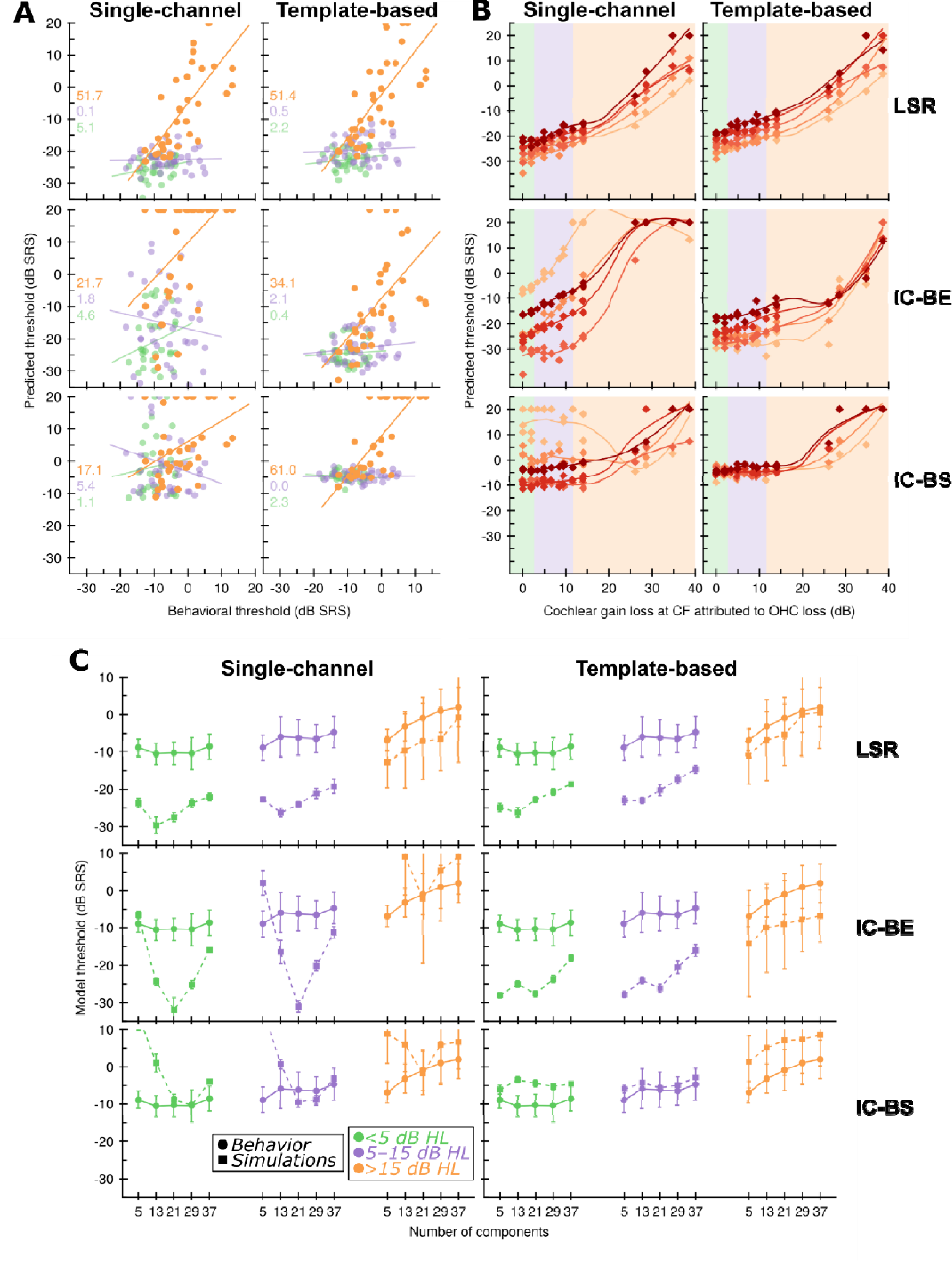
Results of simulations with impaired peripheral function. A. Correlations between behavioral profile-analysis thresholds (x-axis) and simulated profile-analysis thresholds (y-axis) for the fixed-level 2-kHz condition for models with simulated audiograms matched to each participant. Here, data were pooled across different component counts. Different observer strategies and model stages are faceted along columns and rows, respectively. Numbers on the left side of each panel indicate the percent variance explained by each regression line. Data from different groups based on audiometric status are indicated with different colors. B. Scatterplots of simulated profile-analysis thresholds (y-axis) versus the COHC model parameter value for the channel tuned closest to the target frequency expressed in terms of cochlear gain loss relative to COHC=1.0 for the fixed-level 2-kHz condition. Different observer strategies and model stages are faceted as in A. The color of markers and lines indicates the component count, with lighter colors corresponding to fewer components. Dashed curves indicate predictions from a LOESS model fit separately to data from each component count. C. Thresholds for the fixed-level 2-kHz conditions averaged across subjects (or simulated subject audiograms) as a function of component count, for either behavioral data (solid lines and circles) or simulations (dashed lines and squares). Error bars indicate +/− 1.96 standard error of the mean. Data from different groups based on audiometric status are indicated with different colors and arranged left to right from least hearing loss to most hearing loss.

The relationship between degree of simulated cochlear impairment (i.e., C_OHC_/C_IHC_) and the resulting model profile-analysis threshold differed between different observer strategies and model stages. To quantify this, we simulated a measure of cochlear gain (see Methods for details) as a function of the C_OHC_ parameter for a 2-kHz model CF. These simulations provided a mapping between C_OHC_ and cochlear gain loss (relative to C_OHC_ = 1.0), which was in turn used to transform each listener’s C_OHC_ value into a “cochlear gain loss” value in dB. In Fig. 9B, profile-analysis thresholds are plotted as a function of this estimated cochlear gain loss to explore differences in how sensorineural hearing loss affected coding in each model stage and observer strategy. For moderate amounts of loss (i.e., cochlear gain loss < 15 dB, C_OHC_ > 0.5), both single-channel and template-based LSR thresholds showed similar gradual increases in profile-analysis thresholds with increasing loss. For more extreme amounts of loss, the monotonic relationship persisted but it accelerated upward, with the most severe amounts of loss resulting in very poor or unmeasurable thresholds. For the BE and BS models, relationships between hearing loss and predicted thresholds were more complicated. For the BE model, single-channel thresholds were affected by hearing loss, but more so for some component counts than for others. Template-based thresholds showed little dependence on hearing loss until values beyond 20-dB estimated cochlear gain loss. For the BS model, template-based thresholds were similar to the BE model, showing little change until gain loss exceeded 20 dB. Single-channel thresholds were also fairly insensitive to hearing loss, except in the 5-component condition, where hearing loss actually *improved* thresholds (it is interesting to note that many listeners with significant hearing loss did indeed perform better in some 5-component conditions, as shown in Figs. 3, 4.).

Figure 9C shows average behavioral and predicted thresholds as a function of component count in the 2-kHz condition, separated by degree of hearing loss. One striking trend that can be noted is that variability between listeners (or simulated audiograms) was systematically underestimated for listeners with less than 15 dB HL and systematically overestimated for listeners with greater than 15 dB HL. That is, for listeners in the normal-hearing range, model results were very similar across different simulated audiograms, whereas behavioral thresholds were more variable (compare green and purple errorbars for squares vs. circles, Fig. 9C). In contrast, for listeners with hearing-impairment, model results were highly variable across different simulated audiograms, whereas behavioral thresholds were less variable (compare orange errorbars for squares vs. circles, Fig. 9C). This modeling result suggests that more work remains to characterize and model potential sources of variability in psychophysical data and how those sources may differ among different populations of listeners.

Qualitatively, both single-channel and template-based decoding of the LSR model succeeded in capturing the transition from the U-shaped to flat curve with respect to component count seen in normal-hearing listeners to the upward slope seen in hearing-impaired listeners. However, the LSR model also grossly overestimated the difference between normal-hearing and hearing-impaired listeners by predicting normal-hearing thresholds that were much better than what was seen in behavior. Single-channel decoding of BE responses did not result in good fits to the data. Template-based decoding of BE responses suffered from the same overprediction of the effects of hearing loss as seen in the LSR results. Single-channel decoding of BS responses did not resemble behavior, but template-based decoding of BS responses did a fairly good job of capturing the roughly flat shape of the data seen in normal-hearing listeners, correctly predicted an upward slant of the data in hearing-impaired listeners, and exhibited a smaller over-prediction of the effects of hearing loss in the data as compared to the LSR and BE models.

## IV. DISCUSSION

In the present study, we measured thresholds for detecting a relative spectral increment in an inharmonic complex tone (profile analysis) across a wide range of conditions and in listeners with and without sensorineural hearing loss. We found that profile-analysis thresholds worsened substantially as the target frequency was increased from 1 to 2 kHz and from 2 to 4 kHz. We also found that sensorineural hearing loss at 2 kHz was associated with elevated profile-analysis thresholds at a target frequency of 2 kHz, particularly for stimuli with many spectral components; for other conditions, the relationship was not reliably observed or was difficult to measure due to poor task performance. Predicting task performance based on the responses of simulated modulation-sensitive IC neurons to our stimuli, we were able to account for overall trends with respect to target frequency and degree of sensorineural hearing loss.

The present data and simulations build directly from the modeling work described in Maxwell et al. (2020). Their study focused on modeling performance in the “up-down” variant of the profile-analysis task, which uses a stimulus consisting of alternating amplified and attenuated components, rather than a single amplified target component. For that variant of the task, a BE model similar to our own captured the basic trends with respect to component count seen in human data from Lentz et al. (1999). Our BE model was similar to that of Maxwell et al. (2020), but it did not generalize to our task, as our BE-model thresholds did not capture the relationship between component count and profile-analysis thresholds. However, our BE model was able to explain elevated thresholds at higher target frequencies, suggesting that the basic modeling framework is sound. Future work should explore whether differences between the studies were due to the different stimuli employed or to differences in certain methods, such as IC model parameters, CF range and sampling, or procedural details (e.g., number of simulated trials, simulation of interstimulus intervals).

It is interesting to compare the present modeling results to relevant work from the literature on how profile-analysis stimuli might be represented in the auditory periphery (Deng et al., 1987; Horst et al., 1987). To our knowledge, physiological responses to single-component increments added to *inharmonic* complex tones (as in the present study) have not been recorded, but insights can nevertheless be gleaned from responses to increments added to *harmonic* complex tones in the AN (Deng et al., 1987; Horst et al., 1987). In these studies, ANFs tend to synchronize most strongly to the F0 when the complex-tone spectrum is flat. When an increment is added to a single component, AN responses begin to be dominated instead by synchronization to the frequency of the incremented component, at least when the component is near the fiber’s CF, resulting in flatter response envelopes. The results, along with related results on vowel coding in AN (Young and Sachs, 1979; Delgutte and Kiang, 1984), are consistent with a strong role for cochlear nonlinearity in shaping responses to both small spectral increments, as in profile-analysis-like stimuli, and more ecological spectral peaks, such as formants. The peripheral model that we employed (Zilany et al., 2014) provides a good account of the relevant phenomena, and thus likely provided a reasonably accurate simulation of AN responses to the profile-analysis stimuli used here. The key results in the physiological data echo our own observations about the potentially important effects of cochlear suppression in shaping responses to profile-analysis stimuli (Fig. 7), and more generally the ubiquity of CF-dependent capture in responses to complex spectra motivates the focus on coding of neural fluctuations.

Several of the present modeling results prompt reevaluation of prior profile-analysis results. In particular, our results call into question the idea that listeners actually perform “profile” analysis; that is, that listeners compare responses across different cochlear channels within a single interval to perform the task (Green, 1983). One piece of evidence used previously to justify the notion that listeners analyze a “profile” is that thresholds remain fairly stable under level roving. As discussed, roving produced a large threshold elevation when using single-channel LSR rates, consistent with this line of reasoning. However, the same threshold elevation was not always seen for the single-channel BE and BS models, and even when threshold elevation was observed, thresholds were often still quite good under the rove. Put another way, the relative stability of thresholds in the face of level roving is consistent with single-channel models at the level of the IC, and it need not indicate that listeners rely on a multichannel analysis of the spectral “profile” of a sound.

Another piece of evidence used in support of the notion of “profile” analysis is that thresholds worsen as component count falls to small values and components become distantly spaced. Conventionally, this trend in thresholds is assumed to be a perceptual consequence of reduced availability of independent estimates of the pedestal level as fewer components are in the stimulus (Bernstein and Green, 1987). However, all of our simulated single-channel results exhibited this same worsening in thresholds for low component counts, indicating that the trend does not necessarily connote the use of a “profile”. For the LSR model, this shape was a byproduct of suprathreshold cochlear suppression. For the BE and BS models, suppression likely played a role, but the trend was further exacerbated because low component counts resulted in primarily off-target-CF responses in the BE and BS models (Fig. 7A) and elicited neural fluctuations that were too fast to be well-encoded in the BE and BS models. Regardless of the details, it is clear that interpretation of psychophysical data can be aided by an accurate computational model of the auditory system. In this case, use of such a model revealed that certain features of profile-analysis data could be “mirages” that disappear after taking into account the influences of peripheral filtering, peripheral transduction, and midbrain modulation tuning.

Another key takeaway from the present results pertains to the “dynamic range problem”. Because HSR fibers are known to saturate at low sound levels, many researchers have theorized that listeners must rely on LSR fibers at moderate-to-high sound levels (Viemeister, 1988; Bharadwaj et al., 2014). Our modeling results demonstrate that this is not a safe assumption. Our BE and BS model neurons were driven by rate-saturated HSR fibers —no information from LSR fibers reaches the midbrain of our model—and yet our BE and BS models achieved good thresholds in many conditions. The sensitivity of the BE and BS models to sound features other than energy (or average rate) expands the range of levels over which they can effectively operate, at least for coding certain features such as a spectral peak, without expanding their dynamic range *per se* (i.e., the span of the sloping section of their pure-tone rate-level functions). Given the relative sparsity of neurons with wide dynamic ranges in the ascending auditory pathway (Semple and Kitzes, 1985; May and Sachs, 1992) and the strong evidence that modulation coding is a key feature of central auditory processing (Joris et al., 2004), these modeling results are consistent with the neural-fluctuation theory of suprathreshold sound coding (Carney, 2018).

The present results have several implications for the broader field of auditory perception. Perhaps most notably, they suggest the need for a reevaluation of other frequency effects reported in the literature. Many other auditory abilities are known to degrade at higher frequencies, such as frequency discrimination (Moore et al., 1973; Moore and Ernst, 2012), F0 discrimination (Lau et al., 2027; Gockel et al., 2020; Guest and Oxenham, 2022), and intensity discrimination (Jesteadt et al., 1977; Florentine et al., 1987). However, it remains a puzzle why some degrade so quickly with increases in frequency (e.g., frequency discrimination) and others less so (e.g., F0 discrimination; Lau et al., 2017). Our results suggest that a perspective centered on the interplay between peripheral filtering, peripheral transduction, and midbrain modulation coding could provide a new unifying framework to understand these effects. Accordingly, future modeling work should probe whether the modeling strategy used here can generalize to predict frequency effects in other tasks.

## ACKNOWLEDGEMENTS

The authors would like to thank undergraduate research assistants C. Evelyn Feld and Kylie Roloson for assistance with data collection and analysis.

